# Early life infection with *Cryptosporidium parvum* induces inflammatory responses to dietary antigens

**DOI:** 10.1101/2025.02.07.637163

**Authors:** Iti Saraav, Wanyi Huang, Jisun Jung, Rui Xu, Yong Fu, Keely McDonald, Lawrence A. Schriefer, Rachel Rodgers, Megan T. Baldridge, Rodney D Newberry, Chyi Hsieh, L. David Sibley

**Affiliations:** Department of Molecular Microbiology, Washington University School of Medicine, St. Louis, MO, USA; Department of Internal Medicine, Division of Rheumatology, Washington University School of Medicine, St. Louis, MO, USA; Department of Internal Medicine, Division of Gastroenterology, Washington University School of Medicine, St. Louis, MO, USA; Department of Medicine, Division of Infectious Diseases, Edison Family Center for Genome Sciences and Systems Biology, Washington University School of Medicine, St Louis, MO, USA

**Keywords:** cryptosporidium, enteric infection, regulatory T cells, oral tolerance, dietary antigen, T_H_1 inflammatory response, dysbiosis, intestinal inflammation

## Abstract

To examine the effect of early-life infection with *Cryptosporidium parvum* on the development of oral tolerance, we developed a low-dose infection model in neonatal mice. *C. parvum* infection in neonatal mice results in immunopathology in the colon. IL-1β released during *C. parvum* infection blocked the formation of colonic goblet cell associated antigen passages, which normally serve as a conduit for antigen uptake and development of peripheral regulatory T cells (pTregs), responsible for long-term oral tolerance. Following infection with *C. parvum*, adoptively transferred OT-II cells, which respond to ovalbumin (ova), developed reduced frequency of Foxp3^+^Rorγt^+^ cells in mesenteric lymph nodes with an expansion of T_H_1-like Tregs in the colon. The altered pTreg profile was accompanied by a strong T_H_1 immune response and robust IgG2c antibody responses to orally administered ova. Our findings suggest that intestinal inflammation and altered pTreg development leads to loss of oral tolerance during early life infection with *C. parvum*.

## Introduction

*Cryptosporidium* is a leading cause of diarrheal-related deaths in children under 2 years of age in developing countries (1–3). Molecular epidemiological studies indicate that human infections are primarily caused by *Cryptosporidium parvum,* which is zoonotic and acquired from farm animals, and by *C. hominis*, which is spread human-to-human (4, 5). *Cryptosporidium* is an intracellular parasite that infects the epithelial cells of the small intestine, leading to histological abnormalities such as villus blunting, crypt hyperplasia, and microbial dysbiosis that result in significant nutritional deficiencies and diarrhea (6–8). Immunity against *Cryptosporidium* infection in mice and humans is mediated by both innate and adaptive immune responses (9–11). The innate immune response shapes the adaptive immune response by presenting antigens via dendritic cells (DC) and producing pro-inflammatory cytokines like interleukin (IL-12), which drive the differentiation of naive T cells into T helper (T_H_1) cells. Interferon gamma (IFN-γ), a key cytokine in limiting *Cryptosporidium* replication, acts as a critical bridge between the two arms of immunity (12–14). However, an overactive T_H_1 response during infection can lead to heightened inflammation, resulting in tissue damage and impaired intestinal function. Young children are most severely affected by *Cryptosporidium* infections, which typically wane by the age of 18 to 24 months (1). However, the lingering effects of malnutrition, along with lower height and weight for their age, can persist for years (15). Remarkably, even prior asymptomatic infections with *Cryptosporidium* species can lead to malnutrition and stunted growth in children (16–18). The molecular mechanisms behind this lasting impact, even after early-life infections wane, remain unknown.

In early life, goblet cells play a key role in maintaining immune homeostasis through a unique anatomical feature called goblet cell-associated antigen passages (GAPs), which provide a conduit for sampling luminal contents (19, 20). In mice, the formation of GAPs is regulated by falling levels of epidermal growth factor present in the mother’s milk. This reduction initiates GAP formation in the colon during day of life 11 to 21, known as the “window of tolerance,” and later in the small intestine (21). During the formation of GAPs in the colon, a specific subset of peripheral regulatory T cells (pTregs) expressing both Foxp3 and Rorγt is established, and these long-lived cells contribute to tolerance to commensals and dietary antigens (22–24). Some pTregs are uniquely educated in early life, and disruption of the intestinal immune homeostasis during this time can lead to a loss of oral tolerance in adulthood (21, 25–27). *Salmonella* infection in adult mice reduces small intestinal GAP formation, resulting in suppressed tolerogenic responses to dietary antigens (28). However, the impact of infection on the development of oral tolerance when an enteric pathogen is encountered in early life is still not well understood. Therefore, we investigated how cryptosporidiosis in early life affects oral tolerance to dietary antigens that normally develop during this period.

Previous studies have demonstrated that neonatal mice are more susceptible to *C. parvum* infection than adults, with the infection predominantly affecting the ileum. However, in younger animals, the infection can also spread to the colon (29–31). Infections typically elicit a strong T_H_1 response, characterized by the recruitment of monocytes, dendritic cells, and cytokines production such as IL-1β, IL-12, IL-18, and IFN-γ, which contribute to the resolution of the infection within 3-4 weeks (32–35). However, the majority of these prior studies used high-dose infection and did not examine the effect on oral tolerance, even though these responses develop during early life. To address this knowledge gap, we developed a low-dose infection neonatal mouse model to mimic natural infection dynamics. Our findings reveal that early life infection with *C. parvum* disrupts GAPs in the colon, alters pTreg development, and promotes strong T_H_1 immune responses to dietary antigens.

## Results

### *C. parvum* infection in neonatal mice results in immunopathology in the colon

Given the limited understanding of the impact of *C. parvum* infection on the colon, we aimed to investigate the effects of *C. parvum* infection in this intestinal region in neonatal mice. To monitor the age-dependent impact of *C. parvum* infection along the intestine, we infected 5-, 12-, or 19-day old mice, and six days later, we measured parasite burdens in the ileum and colon of infected mice by qPCR. We observed that the parasite burden was highest in the ileum of 5-day old mice as compared to mice that were older when infected (Fig 1A). Interestingly, infection also extended to the colon, where it was found at substantial levels (Fig 1A). The infection intensity decreased in both the ileum and colon in an age-dependent manner (Fig 1A). At six days post-infection (dpi), the parasite burden peaked in the ileum and colon of mice that were 5-day old when infected, so we chose this time point for all our subsequent studies (Fig S1A). Histological studies confirmed the presence of the parasite in the ileum and colon of infected mice, demonstrating that PCR results did not simply represent residual DNA present in the gut (Fig 1B). To study the extent of early life infection on parasite shedding, we infected 5-day old mice with *C. parvum* and measured parasite burdens from fecal material collected between day 5 to day 28 post-infection. We observed that mice become increasingly resistant to *C. parvum* colonization as they aged (Fig 1C). The lowest levels of parasite shedding in these mice was observed at day 28 post-infection (Fig 1C).

**Figure 1.**
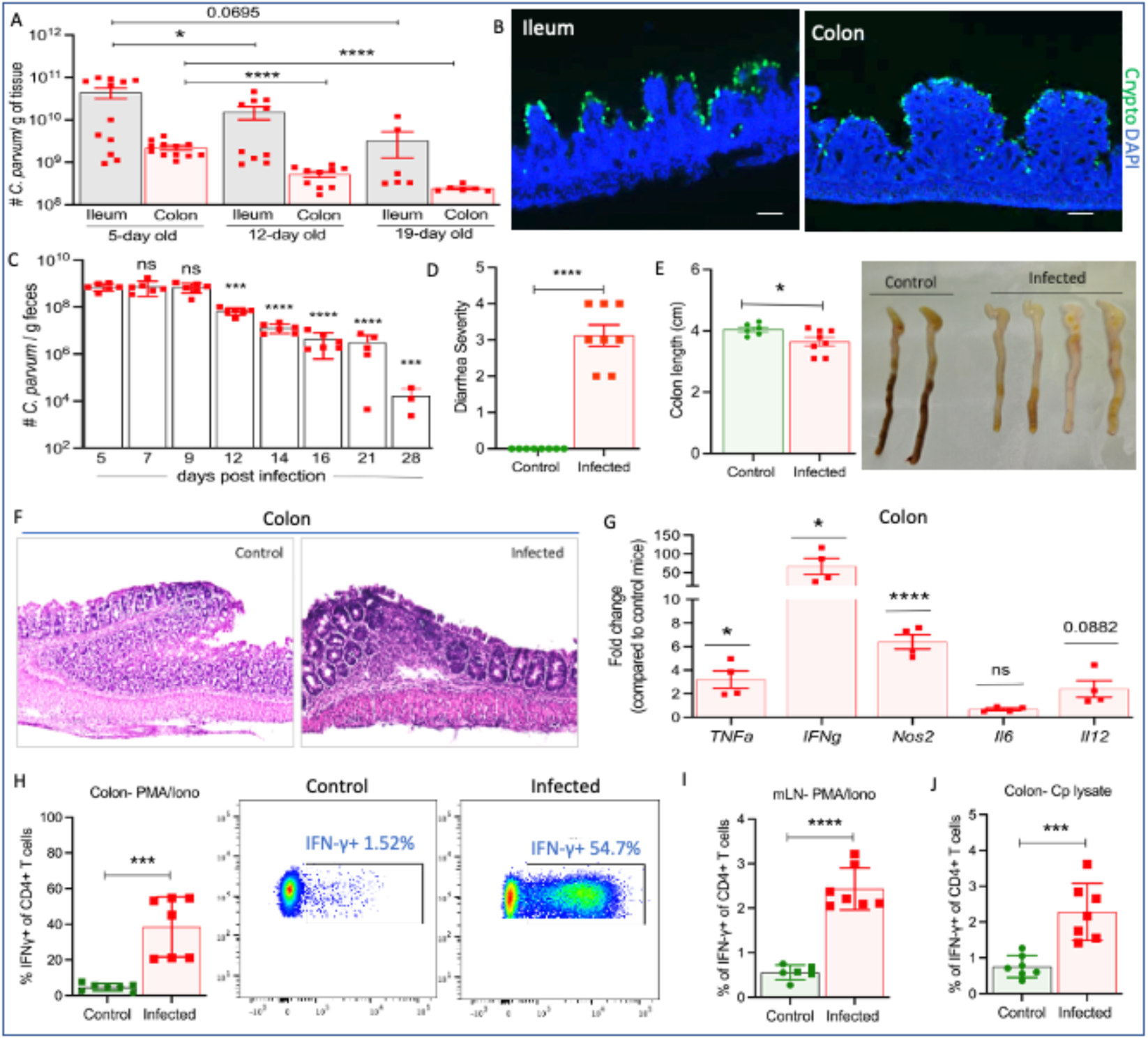
*C. parvum* infection in neonatal mice results in immunopathology in the colon. (A) The average number of *C. parvum* per g of tissue in 5-, 12-, and 19-day old mice monitored 6 days post-infection by qPCR. (B) Sections from the ileum and colon of 5-day old mice examined 6 days post-infection. Parasites stained with rabbit anti-Cp antibody (green), nuclei stained with DAPI (blue). Scale bar, 50 µm. (C) The average number of *C. parvum* per g of feces in 5-day old mice monitored at the indicated days post-infection by qPCR. For statistics, time points were compared to day 5 post-infection. (D) Diarrhea score in 5-day-old mice monitored at day 6 post-infection. Scoring scheme: 1- wet stools, 2- pasty stools, 3- semiliquid, 4- watery diarrhea. (E) Colon length of 5-day old mice measured at day 6 post-infection. (F) Representative image of H&E staining of a section of the colon from 5-day old mice examined at 6 days post-infection. (G) qRT-PCR analysis of mRNA levels of indicated cytokines in the colon of 5-day old mice measured at day 6 post-infection. The frequency of CD4^+^IFN-γ^+^ T cells in (H) colon (I) mLN of 5-day old mice examined at 6 days post-infection obtained after activation with PMA/ionomycin for 3.5 hours and analyzed by flow cytometry. (J) The frequency of CD4^+^IFN-γ^+^ T cells in the colon of 5-day old mice examined at 6 days post-infection obtained after stimulation with *C. parvum* lysate antigen for six hours and analyzed by flow cytometry. (A-F, H-J) Means ± SD from two independent experiments with n = 3-4 mice per group in each experiment. (G) Means ± SD from two independent experiments with n = 2 mice per group compared to average values of control samples from the colon. Statistical tests performed: (A-F, H-J) Unpaired Student’s t-test was used for comparison. (G) One-way ANOVA with Sidak’s multiple comparison. \**P* < 0.05, \*\*\**P* < 0.001, \*\*\*\**P* < 0.0001, ns, not significant. See also Fig S1

To test the effect of *C. parvum* infection on colon inflammation during early life, we infected 5-day old mice. After six days, control and infected mice were monitored for diarrhea, and following euthanasia, colon lengths were measured, and histopathological studies were performed. Infection of neonatal mice was associated with diarrhea (Fig 1D). Additionally, infected mice showed a significant decrease in the length of the colon compared to control mice (Fig 1E). Furthermore, a histological examination of the colon in the infected mice revealed intestinal damage, including cellular infiltration and destruction of the crypt structure, suggesting *C. parvum* infection causes pathology in the colon of neonatal mice (Fig 1F). However, we did not observe any decrease in the body weight of 5-day old mice monitored from day 5 to day 28 post-infection compared to control mice (Fig S1B), indicating this low-dose challenge model has minimal impact on growth.

To determine which cytokines were associated with enhanced colon inflammation during *C. parvum* infection, we used qRT-PCR to analyze the expression of pro-inflammatory genes in the colon of infected mice. We observed a significant increase in the expression of genes such as Tumor necrosis factor (*Tnfa)* and Nitric oxide Synthase *(Nos2)* in the colon of infected mice (Fig 1G). Importantly, there was a highly significant 50-fold increase in the expression of *Ifng* in the colon of infected mice relative to control mice (Fig 1G). To characterize the association of T_H_1 immune responses with gut pathology during *C. parvum* infection, we investigated the T_H_1 phenotype in the CD4 T cell population present in the colon of control and infected mice by flow cytometry. 5-day old mice were infected with *C. parvum,* and six days later, CD4 T cells isolated from the colon lamina propria were analyzed. The colons of infected mice harbored a significantly higher frequency of IFN-γ^+^ producing CD4 T cells as compared with that of control mice (Fig 1H). Importantly, the mesenteric lymph node (mLN) and spleen of infected mice also had a significantly higher frequency of IFN-γ^+^ CD4 T cells (Fig 1I, Fig S1C). To identify the frequency of *C. parvum* specific T_H_1 cells in the colon following infection, 5-day old mice were infected with *C. parvum,* and six days later, cells from colon lamina propria were stimulated *ex vivo* with *C. parvum* lysate antigens. Our flow cytometry data revealed a significant increase in the frequency of IFN-γ^+^ CD4 T cells in the colon of infected mice (Fig 1J). Together, these findings suggest that early life infection with *C. parvum* triggers inflammatory responses that lead to immunopathology in the colon.

### *C. parvum* infection in neonatal mice results in microbial dysbiosis and selective disruption of the gut barrier

We next examined whether *C. parvum* induced effects in the colon of neonatal mice and susceptibility to infection could be due to changes in the microbial composition. To identify changes in bacterial communities, we performed 16S rRNA gene sequencing and analysis of amplicon sequence variants from cecal contents from 5-, 12-, and 19-day old mice sampled at day 6 post-infection (Fig 2A). Infection of 5-day old mice with *C. parvum* resulted in a significant difference in overall bacterial community composition as determined by beta diversity analysis as well as in alpha diversity (Shannon index) relative to control mice (Fig 2B, C). These changes were driven by an expansion of *Enterococcaceae* taxa in mice that were 5-day old when infected but not in mice that were older when infected (Fig 2D, S2A).

**Figure 2.**
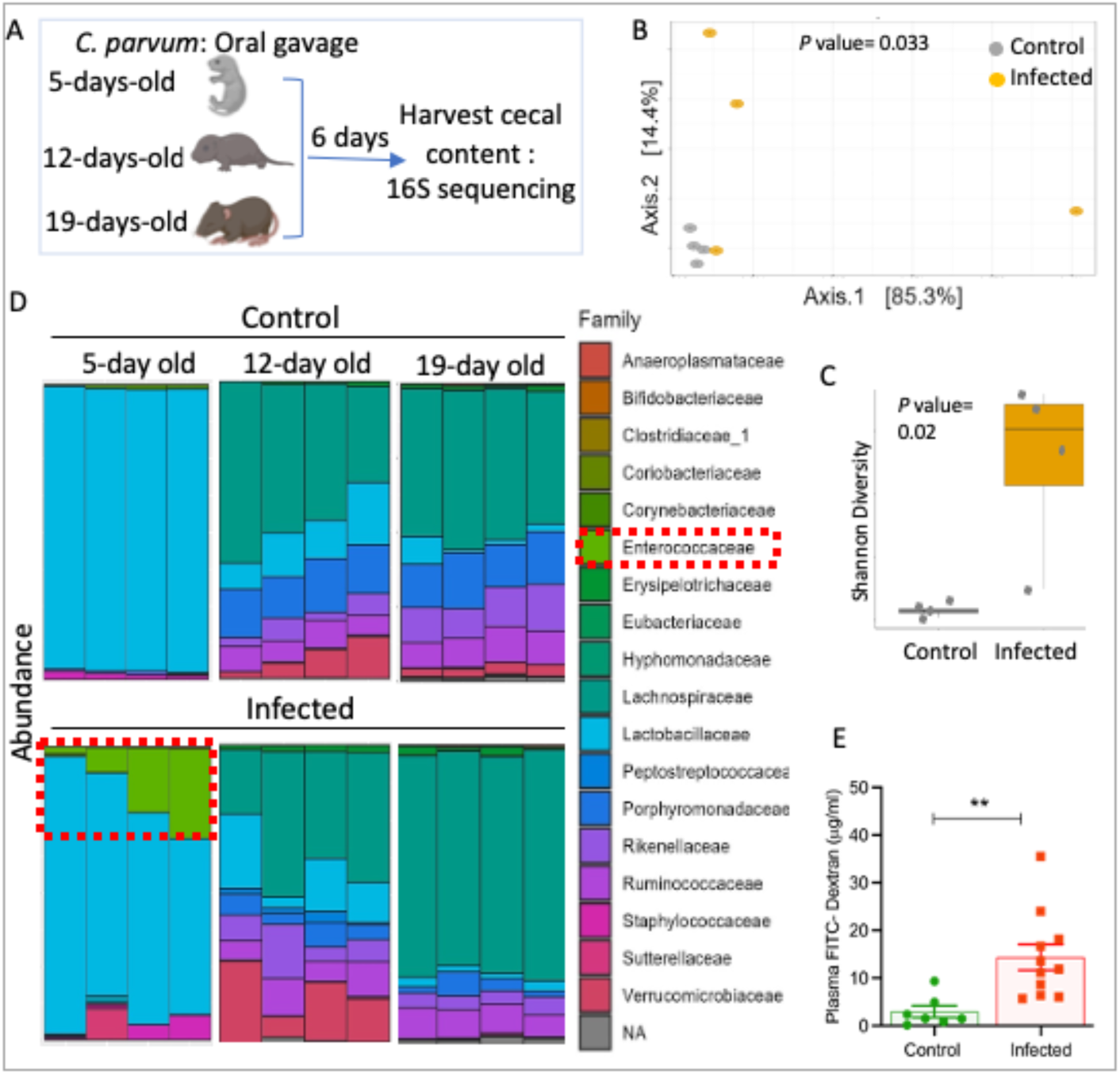
*C. parvum* infection in neonatal mice results in microbial dysbiosis and selective disruption of the gut barrier. (A) Experimental design for collecting samples for microbiota analysis for control and infected mice determined by 16S v4 region rRNA sequencing for Fig B-D. (B) Beta (C) Alpha diversity plots in 5-day old mice analyzed at day 6 post-infection (D) Family level composition in 5-day, 12-day, 19-day old mice sampled at day 6 post-infection (E) The concentration of FITC-dextran in the serum of 5-day old mice measured at day 6 post-infection after 4 h of oral administration of 150μl of 80mg/ml 4kDa FITC dextran. (B, C, E) Means ± SD from two independent experiments with n = 3-4 mice per group in each experiment. Statistical tests performed: (B-C) Kruskal-Wallis, (E) Unpaired Student’s t-test was used for comparison. \*\**P* < 0.01. See also Fig S2

Given the immaturity of the gut barrier in neonatal mice, dysbiosis due to infection could disrupt tight junctions, leading to barrier dysfunction. We further investigated the effect of *C. parvum* infection on intestinal permeability in neonatal mice using a dextran permeability assay. To test this, we infected 5-day old mice, and 6 days later, fluorescein isothiocyanate dextran (FITC-dextran) was administered orally. On average, 12-fold higher levels of FITC-dextran were observed in the serum of the infected mice compared to control mice, suggesting increased intestinal permeability (Fig 2E). We next questioned whether an increase in intestinal permeability upon *C. parvum* infection further results in the translocation of bacteria or bacterial products, such as lipopolysaccharides (LPS). To test this possibility, we infected 5-day old mice with *C. parvum,* and six days later, distal tissues such as mLN and spleen from control and infected mice were harvested and homogenized. Plating of homogenized tissues isolated from control and infected mice on MacConkey and tryptic soy agar plates for 48 h did not result in the formation of any bacterial colonies. Serum analysis of infected mice also revealed normal endotoxin levels similar to control mice (Fig S2B). This result suggests that *C. parvum* infection in neonatal mice leads to paracellular leak where only small molecules can pass through, while larger particles like bacteria and LPS with a molecular weight of about 10–20 kDa are excluded. Together, these findings suggest that *C. parvum* infection in neonatal mice alters the microbial composition but does not lead to bacterial translocation or systemic endotoxemia.

### IL-1β release during *C. parvum* infection in neonatal mice inhibits colonic GAPs

We next evaluated a potential role for *C. parvum* infection in disrupting the formation of GAPs in the colon, which normally form early in life and play a critical role in establishing long-term oral tolerance to food antigens and commensals (36). To test the impact of *C. parvum* infection on GAPs, we infected 5-day old mice, and six days later, 10 kDa Tetramethyl rhodamine (TMR)-labeled dextran was injected into an intestinal loop, mice were euthanized after 25 mins, colonic tissues were processed for histology, and GAPs were visualized and quantified by fluorescence microscopy (Fig 3A). The number of GAPs in the colon of *C. parvum* infected mice was greatly reduced compared to control mice (Fig 3B). Closure of GAPs required live infection and was not induced by the administration of heat-inactivated oocysts (Fig 3B). However, the number of goblet cells was not altered in the infected mice, suggesting that *C. parvum* inhibits GAP formation independent of changes in goblet cell numbers (Fig S3A). Activation of MyD88 signaling through receptors such as Toll-like Receptors (TLRs) or IL-1R on goblet cells can inhibit the formation of GAPs (28, 37). Although *C. parvum* does not cross the epithelial layer, we reasoned its ability to induce inflammatory mediators such as IL-1β and IL-18 in the colon of neonatal mice that may be responsible for the closure of GAPs. To examine this possibility, 5-day old mice were infected, and six days later, supernatant from whole tissue lysates of colons from control and infected mice were analyzed for cytokine production. Strong induction of IL-1β was observed in the colon of infected mice compared to control mice, whereas IL-18 levels were elevated in both conditions (Fig 3C, D). *C. parvum* infection also inhibited GAP formation in the small intestine of adult mice when analyzed at six days post-infection (Fig S3B, C), and analysis of the cytokine production again revealed higher levels of IL-1β in response to infection (Fig S3B, D, E). IL-18 production by epithelial cells has been reported following infection with *C. tyzzeri*, a natural mouse infecting strain (32). To ensure the lack of increased IL-18 production during *C. parvum* infection was not due to differences in cytokine detection methods, we employed the same secretion assay used in the *C. tyzzeri* study for consistency. Sections of the small intestine were removed from adult control and infected mice, and these intestinal explants were assayed for secretion of cytokines by collecting supernatant. Consistent with our previous finding, we observed an increase in the IL-1β levels and not IL-18 during infection (Fig S3F, G). Importantly, the induction of IL-1β required live infection and was not induced by the administration of heat-inactivated oocysts (Fig S3F, G).

**Figure 3.**
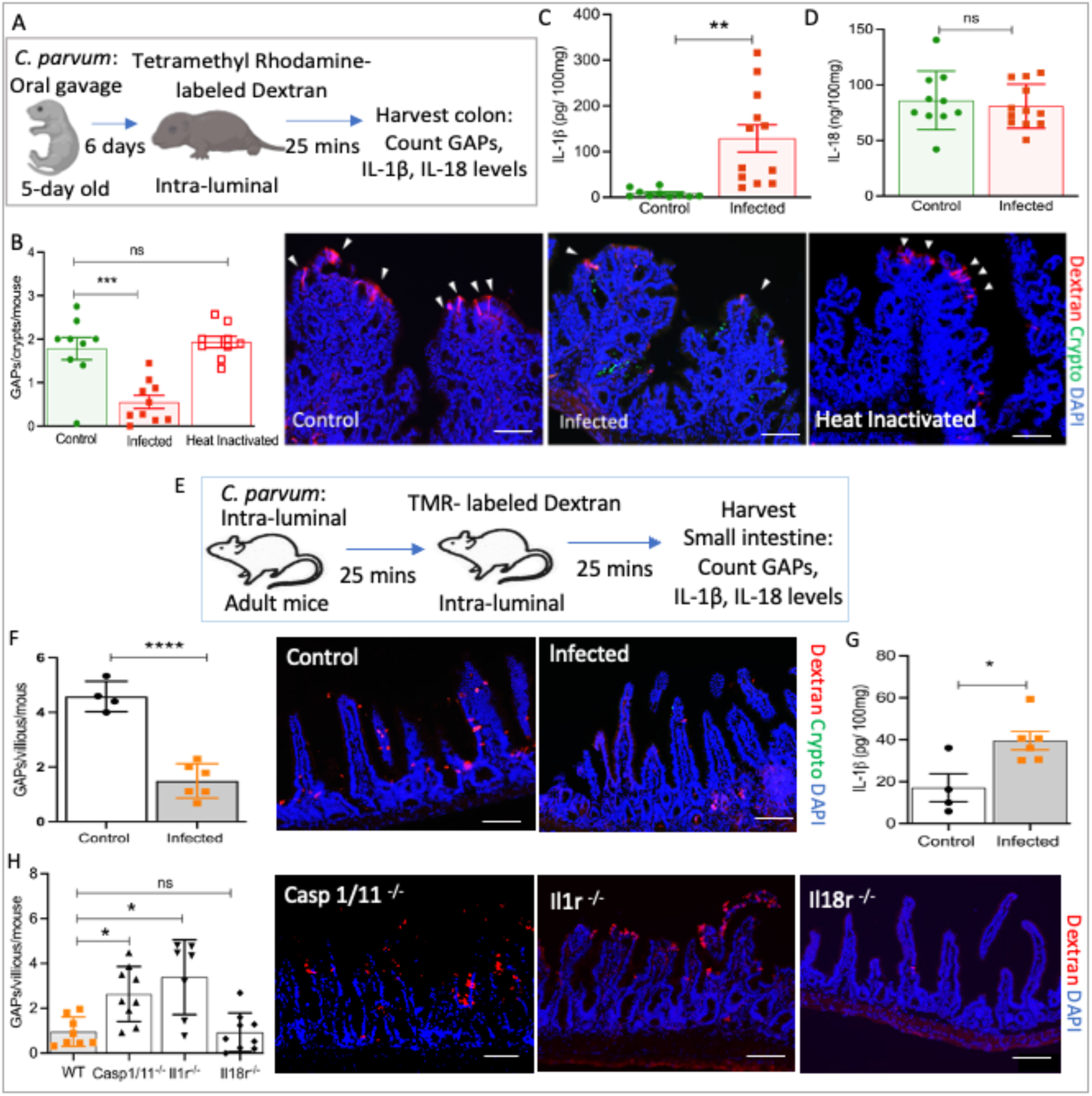
IL-1β release during *C. parvum* infection in neonatal mice inhibits colonic GAPs. (A) Experimental design for counting GAPs and measuring levels of IL-1β and IL-18 in neonatal mice for Fig B-D. (B) Frequency of GAPs in the colon of 5-day old mice infected with *C. parvum* or heat-inactivated oocysts and measured at day 6 post-infection. Representative images are shown with GAPs marked with arrows, Scale bar, 50 µm. Levels of (C) IL-1β (D) IL-18 in whole tissue lysate of the colon of 5-day old mice measured at day 6 post-infection by cytokine bead array. (E) Experimental design for counting GAPs and measuring levels of IL-1β and IL-18 in adult mice infected with *C. parvum* oocysts intraluminally for Fig F-G. (F) Frequency of GAPs in the ileum of control and infected adult mice, and their representative images. (G) Levels of IL-1β in the epithelium scraping of control and infected adult mice by cytokine bead array. (H) Frequency of GAPs in the ileum of indicated knockout mice infected with *C. parvum* and measured at day 6 post-infection, and their representative images, Scale bar, 20 µm. (B-D, H) Means ± SD from two independent experiments with n = 3-4 mice per group in each experiment. (F-G) Means ± SD from two independent experiments, n= 2-3 mice per group in each experiment. Statistical tests performed: (B, H) One-way ANOVA with Sidak’s multiple-comparison test. (C, D, F, G) Unpaired Student’s t-test was used for comparison. \**P* < 0.05, \*\**P* < 0.01, \*\*\**P* < 0.001, \*\*\*\**P* < 0.0001, ns, not significant. See also Fig S3

Previous work has shown that colonic GAPs from preweaning mice are inhibited by heat-killed cecal contents from adult specific pathogen-free (SPF) mice or LPS, suggesting that preweaning mice can respond to microbial products (22, 38). Since our data indicated that *C. parvum* infection in 5-day old mice results in microbial dysbiosis, we addressed whether the altered microbiota due to infection could lead to the closure of GAPs. To test this possibility, we employed an alternative GAP counting approach using direct intraluminal injection into intestinal loops of adult mice, followed by a second injection of TMR labeled dextran, as shown in Fig 3E (28, 39). Using this rapid response assay, we found that intraluminal injection of *C. parvum* oocysts led to the disruption of GAPs and resulted in IL-1β production (Fig 3F, G), suggesting that altered microbiota is not necessary for the closure of GAPs.

To further determine the role of IL-1β and IL-18 in altering the formation of GAPs in the small intestine of mice, we used mice lacking Caspase 1 and 11 (*Casp1/11^-/-^*), the IL-1 receptor (*Il1r1^-/-^*), or the IL-18 receptor (*Il18r^-/-^*). Following the intraluminal injection of *C. parvum* oocysts, we used a similar approach as shown in Fig 3E to count GAPs. Infection of adult wild type or *Il18r^-/-^* mice with *C. parvum* led to inhibition of GAP formation in the small intestine (Fig 3H). In contrast, infection of adult *Casp1/11^-/-^* or *Il1r1^-/-^* mice no longer blocked the formation of GAPs, suggesting the potential role of IL-1β in the closure of GAPs (Fig 3H). Collectively, we conclude that the inhibition of GAP formation during *C. parvum* infection occurs primarily through parasite induction of IL-1β rather than being dependent on microbiota-induced effects.

### *C. parvum* infection in neonatal mice reduces pTregs and increases T_H_1 inflammatory response to dietary antigens in the mLN

We next examined whether the closure of GAPs due to *C. parvum* infection altered pTregs specific to dietary antigens in neonatal mice. To test this possibility, 5-day old mice were infected, and 6 days later, mice received ovalbumin (ova) orally one time and in the drinking water provided to dams for seven days. The next day, cell trace violet (CTV)-labeled ova-specific OT-II T cells were adoptively transferred into the pups. CD4 T cell responses to luminal ova were then monitored by observing the fate of the transferred naive OT-II T cells in the mLN of control and infected mice (Fig 4A). Following one week of ova administration, infection resulted in an enlarged mLN due to an influx of immune cells. We observed an increased number of OT-II T cells in the mLN of infected mice compared to control mice (Fig 4B), with no difference in the number of CD4 T cells (Fig S4B). OT-II cells from both control and infected mice in the mLN proliferated well in response to luminal ova, indicating ova-specific CD4 T-cell activation (Fig 4C). However, a significantly higher frequency of proliferating OT-II T cells was observed in the mLN of infected mice (Fig 4C). *C. parvum* infection significantly reduced the frequency of ova-specific Foxp3^+^Rorγt^+^ Tregs in the mLN as compared to control mice but did not have a significant impact on their absolute numbers (Fig 4D, Fig S4C). Surprisingly, *C. parvum* infection did not significantly reduce the frequency or number of ova-specific Foxp3^+^ Tregs (Fig 4E, Fig S4D). We next addressed whether ova-specific T cells would differentiate to become effector cells during infection. Strikingly, upon transfer into *C. parvum* infected hosts, OT-II T cells differentiated toward a T_H_1 phenotype, as demonstrated by the increase in the frequency and number of OT-II T cells expressing the transcription factor T-bet (Fig 4F, Fig S4E). Analysis of recipient polyclonal T cells after *C. parvum* infection revealed decreased frequencies of Foxp3^+^Rorγt^+^ Tregs in the mLN and an increase in the frequency and number of T-bet^+^ T_H_1 cells (Fig S4H, S4J, S4K). These findings indicate that infection with *C. parvum* in neonatal mice skews CD4 T cell responses toward a T_H_1 inflammatory phenotype and reduces the development of pTregs specific to dietary antigens.

**Figure 4.**
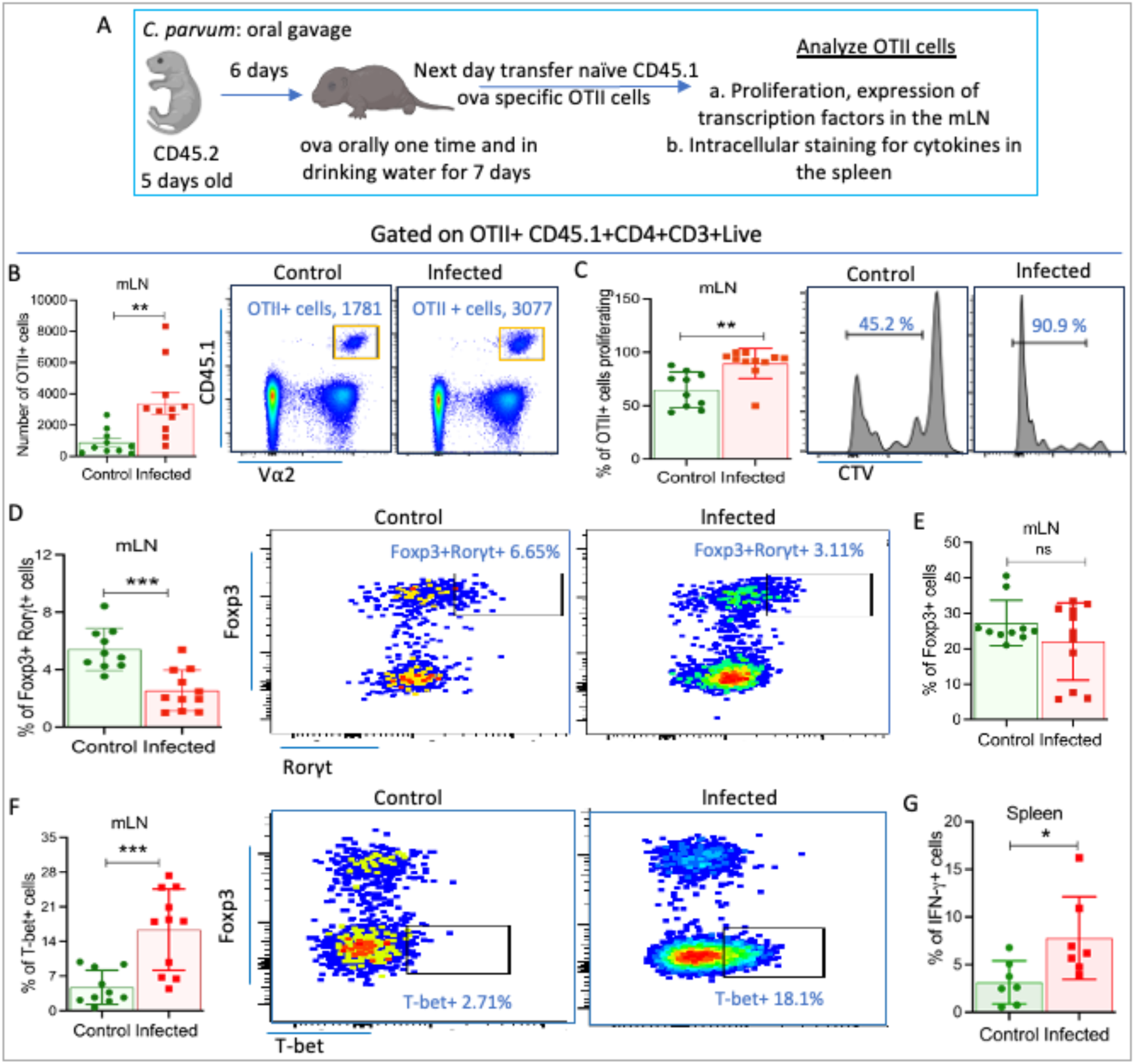
*C. parvum* infection in neonatal mice reduces pTregs and increases T_H_1 inflammatory response to dietary antigens in the mLN. (A) Experimental design for monitoring CD4 T cells response to dietary antigens in the mLN after *C. parvum* infection, ovalbumin (ova) (B) Number of OT-II cells in the mLN of control and infected mice. Frequency of (C) ova-specific proliferating cells (D-F) different transcription factor-expressing ova-specific cells in the mLN of control and infected mice as shown. (G) Frequency of ova-specific IFN-γ^+^ cells in the spleen of control and infected mice. (B-G) Means ± SD from two independent experiments, n= 3-4 mice per group in each experiment. Statistical tests performed: Unpaired Student’s t-test, \**P* < 0.05, \*\**P* < 0.01, \*\*\**P* < 0.001, ns, not significant. See also Fig S4

Ova-specific Tregs can migrate into the spleen and to the peripheral tissues, where they help suppress systemic immune responses (40, 41). Following one week of ova administration, activated OT-II T cells were detected equally in the spleen of both control and infected mice (Fig S4L). The absence of increased OT-II T cell accumulation in the spleen during infection suggests that OT-II T cells preferentially migrate to the mLN to initiate immune responses to *C. parvum* (Fig 4B, Fig S4L). Furthermore, we observed a significant increase in the frequency of IFN-γ^+^ OT-II T cells in the spleen of infected mice (Fig 4G, Fig S4M). Taken together, our data show that during *C. parvum* infection, CD4 T cell tolerance towards dietary antigens is lost, and ova-specific T cells acquire an effector phenotype and respond in a manner comparable with pathogen-specific T cells.

### *C. parvum* infection in neonatal mice expands T_H_1-like Tregs and increases T_H_1 inflammatory response to dietary antigens in the colon

We next investigated the effect of early-life infection on the development of pTregs to dietary antigens in the colon, an important secondary site of infection in neonatal mice. To assess the status of pTregs in the colon, we monitored CD4 T cell responses to luminal ova by tracking the fate of transferred naive OT-II T cells in the colon of both control and infected mice (Fig 5A). Ova-specific T cells were not detected in the colon on day 7 of ova administration but were found in the colon on day 14 (Fig 5B), suggesting that ova-specific T cells are activated in the mLN by day 7 and migrate to the colon by day 14. Consistent with the results observed in the mLN after infection, we noticed an increased accumulation of OT-II T cells in the colon compared to control mice (Fig 5B). We also observed a significant increase in the frequency and number of CD4 T cells in the colon of infected mice (Fig S5B, C). Surprisingly, in *C. parvum* infected mice, OT-II pTreg cell differentiation was unaffected in the colon, and a similar frequency of ova-specific Foxp3^+^ Tregs and ova-specific Foxp3^+^Rorγt^+^ Tregs was observed in control and infected mice upon ova administration (Fig S5D, E). Interestingly, analysis of transcription factors in the colon revealed that pTregs acquire a T_H_1-like Treg phenotype upon entering the colon of infected mice, as shown by the expression of the transcription factor T-bet in pTregs. We observed an increase in the frequency and number of ova-specific Foxp3^+^T-bet^+^, ova-specific Foxp3^+^Rorγt^+^T-bet^+^ Tregs, and ova-specific T-bet^+^ T_H_1 cells in the colon of the infected mice (Fig 5C-H). Analysis of colon polyclonal T cells revealed a significant decrease in the frequency of Foxp3^+^, and Foxp3^+^Rorγt^+^ Tregs and an increased frequency and number of T-bet^+^ T_H_1 cells in the colon after *C. parvum* infection (Fig S5F, S5H, S5J, S5K). Overall, the findings provide evidence that *C. parvum* affects the pTreg population specific to dietary ova, and ova-specific T cells acquire a unique T_H_1-like Treg phenotype in the colon.

**Figure 5.**
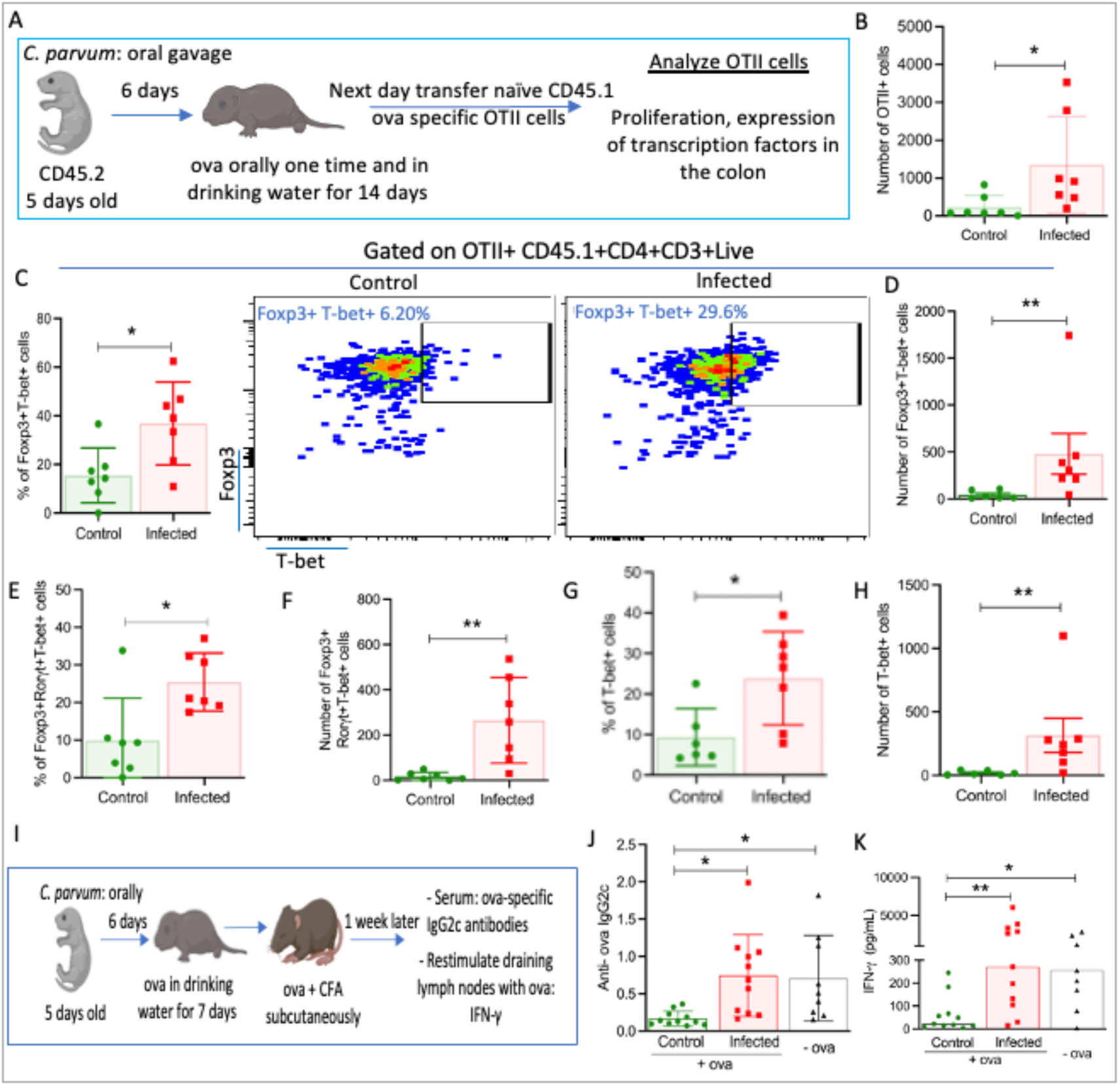
*C. parvum* infection in neonatal mice expands T_H_1-like Tregs and increases T_H_1 inflammatory response to dietary antigens in the colon. (A) Experimental design for monitoring CD4 T cells response to *C. parvum* infection in response to dietary antigens in the colon. (B) Number of OT-II cells in the colon of control and infected mice. (C-H) The frequency or number of different transcription factor-expressing ova-specific cells in the colon of control and infected mice as shown. (I) Experimental design of assay to measure the immune response to ova for Fig J-K, complete freunds adjuvant (CFA) (J) Levels of ova-specific IgG2c antibodies in the serum as measured by ELISA. (K) Levels of IFN-γ in the supernatant of cultured draining lymph nodes isolated from control and infected mice restimulated with ova and quantified by cytokine bead array (B-H) Means ± SD from two independent experiments, n= 3-4 mice per group in each experiment. (J) Means ± SD from two independent experiments, n= 5-7 mice per group in each experiment. (K) Median from two independent experiments, n= 5-7 mice per group in each experiment. Statistical tests performed: (B, C, E-G) Unpaired Student’s t-test, (D, H) Mann-Whitney test (J, K) One-way ANOVA with Sidak’s multiple-comparison test. \**P* < 0.05, \*\**P* < 0.01, ns, not significant. See also Fig S5

To assess the impact of *C. parvum* infection on the immune response to dietary ova, we conducted a functional assay to evaluate the ability of mice to mount a peripheral inflammatory response to an antigen for which immune tolerance had been induced through oral administration (Fig 5I). Infected mice developed a robust IgG2c antibody response to ova, indicating that *C. parvum* induced a T_H_1 immune response (Fig 5J). Subsequently, draining lymph nodes from ova-fed infected mice that were restimulated with ova in vitro showed an increase in IFN-γ secretion compared to control mice (Fig 5K). Therefore, our data support that *C. parvum* infection in neonatal mice alters the development of pTregs to dietary antigens and instead results in a heightened inflammatory immune response to them.

## Discussion

Early life infection with *C. parvum* or C*. hominis* can predispose children to long-term growth stunting and delayed development, although the mechanisms driving these lasting effects are unknown. The establishment of oral tolerance is an important early life event, and perturbation of this process can have long-lasting effects, including impaired tolerance to dietary antigens and an increased risk of food allergies (21). To test the impact of *C. parvum* infection on oral tolerance to food antigens, we used a low-dose infection neonatal mouse model. Our findings demonstrate that infection of neonatal mice with *C. parvum* infection leads to immunopathology, including increased IL-1β production, and leads to closure GAPs in the colon. The closure of GAPs is associated with an altered response to dietary antigens. Specifically, *C. parvum* infection reduced the development of the pTregs to orally administered ova while expanding CD4 T cell responses toward a T_H_1 inflammatory phenotype. Together, our work reveals that early-life infection with *C. parvum* triggers a heightened immune response to dietary antigens, driving inflammation that potentially leads to lasting impairment.

Neonatal individuals, including mice, calves, and humans, are particularly susceptible to *C. parvum* and *C. hominis* infection (42, 43). Importantly, certain immune mechanisms are uniquely developed in the colon during early life, and enteric infection during this time has long-term consequences (21, 44). Here, we employed a low-dose infection neonatal mouse model to investigate the effects of *C. parvum* infection during early life. Our model provides insights into the age-dependent dynamics and impact of *C. parvum* infection in neonatal mice. Although the ileum was the primary site of parasite colonization, substantial infection in the colon was also observed, peaking at six days post-infection in 5-day old mice. Consistent with prior work, parasite burden and shedding decreased with age, reflecting increased resistance to colonization (45). This age-related resistance underscores the importance of studying early-life infections to understand susceptibilities that may not be manifest in adulthood. Infection of neonatal mice with *C. parvum* was associated with significant intestinal damage in the colon, including histological abnormalities in the crypt, and reduced colon length, resulting in diarrhea. Notably, these outcomes occurred without affecting body weight, likely due to the lower *C. parvum* dose used in the present study. This contrasts with earlier studies that reported more pronounced weight loss using higher inoculum (31, 34). We observed a marked upregulation of pro-inflammatory cytokines in the colon of infected mice, including an increased frequency of IFN-γ^+^ CD4 T cells specific to *C. parvum* antigens. Studies in the ileum of adult and neonatal mice have shown that IFN-γ is important for controlling *Cryptosporidium* infection (14). IFN-γ is initially produced by innate lymphocyte cells (ILC1) and intraepithelial lymphocytes for a rapid immune response, while T_H_1 cells become the dominant source during the adaptive immune response (12–14, 46). We recently characterized a strain of *C. tyzzeri* named Ct-STL that behaves as a non-pathogenic commensal in immunocompetent mice (47). Interestingly, Ct-STL colonized various regions of the gastrointestinal tract and elicited immunological effects in the colon, inducing strong T_H_1 responses and elevated IL-12 and RANTES levels (47). However, studies on *Toxoplasma gondii and C. tyzzeri* suggests that an overactive T_H_1 response can lead to tissue damage and makes mice more susceptible to DSS-mediated colitis (47–49). In line with this, we observed that *C. parvum* infection in neonatal mice induces T_H_1-mediated immunopathology in the colon.

Our study reveals that early life infection with *C. parvum* inhibits GAP formation in the colon and emphasizes the necessity of active parasite invasion for this immune modulation. Previous studies have shown that colonic GAPs close during development due to the activation of MyD88 signaling pathways through receptors such as toll like receptors (TLRs) in response to elevated bacterial levels that greatly expand in early life (22). Therefore, we considered whether the expansion of certain taxa during enteric infection with *C. parvum* may have led to GAP closure by acting directly on TLRs. However, direct luminal injection of *C. parvum* oocysts into adult mice also induced rapid GAP closure, indicating that this response occurs independently of microbiota changes. Instead, we confirmed the release of IL-1β during *C. parvum* infection inhibits colonic GAPs, suggesting that parasite-induced cytokine responses are responsible for the disruption of GAPs. Both IL-1β and IL-18 signal through MyD88, with IL-1β previously shown to disrupt small intestinal GAPs during *Salmonella typhimurium* infection (28). Previous studies have demonstrated that IL-18 is strongly induced in the ileum following infection with *C. tyzzeri* (32). In contrast, IL-18 production was not significantly elevated in our model of low-dose *C. parvum* infection, consistent with another study of neonatal mice infected with *C. parvum* (34). Moreover, the loss of the IL-18 receptor did not alter the disruption of GAPs following *C. parvum* infection. This lack of IL-18 induction compared to *C. tyzzeri* infection may reflect strain-specific differences in how *Cryptosporidium* interacts with host cells, potentially activating distinct inflammasome pathways and cytokine responses. Previous work in adult mice has demonstrated that the closure of GAPs during *Salmonella* infection is a physiologically beneficial mechanism for limiting pathogen dissemination (28). In contrast, *C. parvum* occupies an extra-cytoplasmic niche at the apex of intestinal epithelial cells and does not require access to the lamina propria for its survival or replication. Hence, GAP closure during *C. parvum* infection is likely a byproduct of the host immune response rather than a direct defense mechanism against the pathogen.

Despite colonic GAPs being closed, we observed an increase in OT-II T cell proliferation in the mLN during *C. parvum* infection in neonatal mice. This response may be due to increased dendritic cell (DC) sampling of antigens, consistent with the increased recruitment of DCs to the site of infection in neonatal mice infected with *C. parvum* (50). In contrast, we did not observe paracellular leakage of large molecules like ova (42.5 kDa), although *C. parvum* infection in neonatal mice did increase leakage to smaller molecules (<4 kDa). Despite the increased expansion of OT-II cells, *C. parvum* infection in neonatal mice significantly impairs the development of ova-specific Foxp3^+^Rorγt^+^ Tregs in the mLN. This impairment, combined with the closure of GAPs, suggests that this pathway is uniquely critical for generating pTregs during early life (21). In contrast, infection of 3-week-old mice with the mouse commensal *C. tyzzeri* did not alter oral tolerance to ova given 3 weeks later (51). Additionally, the inhibition of GAPs during *Salmonella* infection in adult mice has been shown to reduce ova-specific Foxp3^+^ Tregs without increasing inflammation or altering responses to dietary antigens fed orally (28). Taken together, these results suggest that the timing and possibly degree of inflammation are important in altering Treg populations.

During *C. parvum* infection, ova-specific pTregs displayed a unique phenotype in the colon characterized by the expression of the T_H_1-associated transcription factor T-bet. Similar T_H_1-like Tregs have been reported in *Toxoplasma*, viral infection, and tumor models, where Tregs adapt to inflammatory signals, maintaining regulatory function while acquiring pro-inflammatory characteristics (52–56). For example, lethal challenge with *T. gondii* leads to loss of Foxp3^+^ Tregs in response to orally administered ova and instead, Tregs transition to T-bet^+^ T_H_1 profile leading to overt pathology (56). Our study expands on this finding by demonstrating that such changes can occur with non-lethal infection and that they are associated with altered antigen sampling due to the closure of GAPs. Hence, even low-grade enteric infections may predispose the immune system to polarize toward the T_H_1 profile and compromise the tolerance of oral antigens with potentially long-term effects. Previous studies with reovirus and norovirus have shown that type I IFN signaling in DCs, mediated by the upregulation of interferon regulatory factor 1, drives the production of cytokines, such as IL-12, that promote T_H_1 responses to dietary antigens (57, 58). Notably, *Cryptosporidium* contains a double-stranded RNA virus, *Cryptosporidium parvum* virus 1 (CSpV1), which could engage similar type I IFN signaling pathways and may alter DC function (59, 60). These findings suggest that in addition to GAP closure, altered DC functions might contribute to an increased T_H_1 inflammatory response to dietary antigens observed during *C. parvum* infection.

Early life infections are increasingly recognized as key modulators of immune homeostasis, with the potential to predispose individuals to inflammatory disorders such as food allergies, inflammatory bowel disease, or autoimmune conditions (61, 62). Recent studies and clinical trials have highlighted the time-dependent benefits of introducing food allergens before or around 6 months of age, significantly reducing the risk of developing food allergies (63, 64). This evidence underscores the importance of the timing of antigen exposure in shaping immune responses during early life. Our work contributes to this growing knowledge by demonstrating how early life infections can reshape CD4 T cell differentiation to dietary antigens in the gut and disrupt oral tolerance. Potentially, these altered immune responses in early life may predispose individuals to long-term consequences even after infections with *Cryptosporidium* are resolved.

## Experimental model and subject details

### Mice

Animal studies were conducted according to the US Public Health Service policy on human care and the use of laboratory animals. Animal breeding and experiments were performed in a specific pathogen-free animal facility using protocols approved by the Washington University Institutional Animal Care and Use Committee. C57BL6 timed pregnant mice (strain number: 000664) were purchased from the Jackson laboratory (Bar Harbor, ME). Male and female mice were used to perform experiments. *Il1r1 ^−/−^, Il18r ^−/−^,* and *Caspase^1/11^* knockout mice were a gift from Dr. Stallings (Washington University in St Louis). CD45.1^+^OT-II^+^ transgenic mice were a gift from Dr. Hsieh (Washington University in St. Louis). OT-II transgenic mice were bred to Rag1^−/−^ and Foxp3^IRES-GFP^. Mice were co-housed with siblings of the same sex throughout the experiments.

## Method details

### *C. parvum* oocysts preparation and mouse infection

*Cryptosporidium* Strain *C. parvum* oocysts (AUCP-1 isolate) were maintained by repeated passage in male Holstein calves and purified from fecal material after sieve filtration, Sheather’s sugar flotation, and discontinuous sucrose density gradient centrifugation as previously described (65). All calf procedures were approved by the Institutional Animal Care and Use Committee (IACUC) at the University of Illinois Urbana-Champaign. Purified oocysts were stored at 4 °C in PBS + 50 mM Tris-10 mM EDTA, pH 7.2 for up to six months before use. Before infection, *C. parvum* oocysts were treated in a 40% bleach solution (commercial bleach containing 8.25% sodium hypochlorite) diluted in phosphate-buffered saline (PBS) (Corning) for 10 mins on ice, then washed three times in PBS containing 1% Bovine Serum Albumin (BSA) (Sigma). Bleached oocysts were stored for up to 1 week in PBS plus 1% BSA at 4 °C before infection. Neonatal mice were orally challenged with 2 × 10^4^ oocysts resuspended in 50 μl PBS, and adult mice were orally challenged with 5 × 10^4^ oocysts resuspended in 100 μl PBS.

### Parasite burden

At a specified time postinfection, the tissue of interest was extracted from each mouse, weighed, and then homogenized using 1.5 mm zirconia silica beads (BioSpec) in the Bead Beater (BioSpec). DNA was extracted using the QIAamp fast DNA stool mini kit (Qiagen) with the modifications as previously described (45). Quantitative PCR (qPCR) for the *C. parvum* 18S rRNA was performed using the following primers: ChvF18S (59-CAATAGCGTATATTAAAGTTGTTGCAGTT-39) and ChvR18S (59-CTGCTTTAAGCACTCTAATT TTCTCAAA-39) (57). For qPCR, each 25 μl reaction contained a final concentration of 100 nM for both forward and reverse primers (IDT, San Diego, California) and 12.5 μl SYBR Green PCR Master Mix (Bio-Rad Laboratories). Genomic DNA (2 μl) was added, and the qPCR was performed in QuantStudio3 (Applied Biosystems). Data was analyzed using QuantStudio Design and Analysis Software (Applied Biosystems). Each sample was run in duplicate. A control with no template was run concurrently and was consistently negative. The number of *C. parvum* genomic equivalents was calculated for each sample based on a standard curve using DNA from known quantities of *C. parvum* oocysts and divided by the original weight of the intestinal sample to obtain the number of *C. parvum* organisms per gram of intestine.

### Parasite lysate preparation

The oocyst stock was vortexed, and 2.5 × 10⁸ oocysts were centrifuged at 2,500 ×g for 5 mins at 4 °C. After discarding the supernatant, the pellet was resuspended in PBS (pH 7.2) with 1X protease inhibitor cocktail (Sigma). Oocyst walls were loosened by five freeze-thaw cycles (2 mins in liquid nitrogen, 5 mins at room temperature) followed by cooling on ice for 10 mins. The oocysts underwent five cycles of ultrasonication on ice (30 sec sonication, 1 min rest) to release proteins. The sample was then cooled on ice for 10 mins and centrifuged at 10,000 ×g for 5 mins at 4 °C to remove debris. The supernatant was filtered using an Amicon Ultra-4 10K centrifugal filter (Merck Millipore). Protein concentration was measured using a Bradford assay, and aliquots were stored at −80 °C until use.

### Quantitative RT-PCR

RNA was extracted from the mouse tissue using the RNeasy Micro kit following the manufacturer’s recommendations (Qiagen). cDNA was synthesized from RNA using an iScript cDNA synthesis kit for RT-PCR (Bio-Rad). Real-time PCR of *Tnfa, Ifng, Nos2, Il1b, Il6, and Il12* was performed using SYBR Green PCR Master Mix (Bio-Rad Laboratories). Data acquisition was done in QuantStudio3 (Applied Biosystems) and analyzed using QuantStudio Design and Analysis Software (Applied Biosystems). The expression of target mRNA was calculated and normalized to the expression of the housekeeping gene β-actin using the 2(-ΔΔCT) method (66). Primers are listed in Table S1.

### Histology

Mouse intestines were obtained, and the luminal contents were flushed with cold PBS. Intestines were cut open longitudinally, pinned out, and fixed in 4% paraformaldehyde (Electron Microscopy Sciences) for 16 h at 4 °C. The fixed tissues were washed three times in PBS, followed by dehydration in 70% ethanol. Tissues were then embedded in 2% agar, followed by embedding in paraffin and sectioned at 5 µm. The obtained sections were stained with hematoxylin and eosin for histological evaluations, including damage in tissue architecture and infiltration of immune cells.

### Immunohistochemistry

To detect the presence of the parasites, mice were euthanized, and the intestinal tissue was removed, flushed with PBS, cut open longitudinally, and pinned out and fixed in 4% paraformaldehyde for 16 h at 4 °C. The fixed tissues were washed three times in PBS followed by 30% sucrose overnight. The tissue was frozen in optimal cutting temperature (OCT) solution (Sakura Finetek) and cryosectioned. For immunostaining, sections were thawed and incubated in PBS for 10 minutes, and antigen retrieval was performed with Trilogy (Sigma-Aldrich). Then, non-specific staining was blocked in the sections using PBS-0.1% Triton-10% bovine serum albumin (BSA) (TSS buffer) for 1 h at room temperature. Tissue sections were stained using primary antibody 1: 2000 PanCp antibody (Polyclonal antibody that labels all major stages of *C. parvum,* (67)) prepared in TSS buffer for 1 h at room temperature in a humidified chamber. Sections were washed three times in TSS buffer and incubated with Goat anti-rabbit Alexa Flour 488 secondary antibody (Thermo Fisher Scientific) diluted 1: 2000 in TSS buffer for 1 h at room temperature. Nuclear DNA was stained with Hoechst (Thermo Fisher Scientific) diluted 1: 2000 in PBS for 10 mins at room temperature. Fluorescent and bright field images were collected on a Zeiss Axio Observer inverted microscope with a color camera. Zen software was used to acquire images.

### Diarrhea score and colon length measurement

Five-day old mice were infected with oocysts by oral gavage. At six days post-infection, stool samples were collected by gently massaging mice. Stool samples were scored using a scheme. In this scheme 1-wet stools, 2-pasty stools, 3-semiliquid, 4-watery diarrhea. Mice were then sacrificed, and the colon length was measured using a centimeter ruler.

### 16S rRNA sequencing and analysis

Cecal content was collected and stored at −80 °C until further processing. DNA was extracted using the Pro fecal DNA extraction kit (Qiagen). Primer selection and PCR amplification were performed as described previously (49, 68). Briefly, each sample was amplified in triplicate with Golay-barcoded primers specific for the V4 region (F515/R806) and checked by gel electrophoresis. The final pooled samples, along with aliquots of the three sequencing primers, were sequenced using a paired-end 250bp Illumina protocol by the Center for Genome Sciences (Washington University School of Medicine). Read quality control and the resolution of amplicon sequence variants (ASVs) were performed using the dada2 R package (69). ASVs that were not assigned to the kingdom Bacteria were filtered out. The remaining reads were assigned taxonomy using the Ribosomal Database Project (RDP trainset 16/release 11.5) 16S rRNA gene sequence database (70). Ecological analyses, such as alpha-diversity (richness, Shannon diversity) and beta-diversity analyses (unweighted and weighted UniFrac distances), were performed using PhyloSeq and additional R packages (71).

### FITC-dextran intestinal permeability assay

To detect paracellular leak, 150 μl of 80 mg/ml 4KDa FITC dextran (Sigma-Aldrich FD4) was gavaged into 5-day old mice. Four hours later, serum was collected, and FITC concentration in the sera was measured using a FITC-dextran standard curve in a black 96-well microplate (Greiner BioOne). Fluorescence was measured at 530 nm with excitation at 485 nm using BioTek Cytation 3 cell imaging multimode reader (Agilent).

### GAPs and Goblet cells staining

Colonic and small intestine GAPs were enumerated on fixed tissue sections as previously described (39). Briefly, tetramethylrhodamine-labeled 10 kD dextran (Thermo Fisher) was administered in the intestinal tissue of interest of anesthetized mice (protocol approved to use a controlled substance). After 25 mins, mice were sacrificed, and tissues were thoroughly washed with cold PBS before fixing in 4% PFA. Tissues were embedded in OCT solution (Fisher Scientific), 7-μm sections were prepared, stained with Hoechst (Thermo Fisher Scientific), and imaged using a Zeiss Axio Observer inverted microscope. GAPs were identified as dextran-filled columns traversing the epithelium and containing a nucleus and were counted as average GAPs/crypt/mouse in the colon or GAPs/villus/mouse in the small intestine. Goblet cells in the intestine were stained using the lectin wheat-germ agglutinin and detected using an Immunofluorescence microscope (Zeiss Axio Observer).

### Cytokines levels measurement

To measure the concentration of IL-1β and IL-18 in the colon of neonatal mice, the colon was weighed and homogenized in PBS using a Bead Beater (BioSpec). Cells were centrifuged, and the supernatant was collected. To measure the concentration of IL-1β and IL-18 in the small intestine epithelium of adult mice, a portion of the ileum was opened longitudinally and washed with PBS. The epithelium cell layer was scraped and weighed, and cells were homogenized in PBS. Cells were centrifuged, and the supernatant was collected. To measure the levels of IL-1β and IL-18 secreted by the intestinal tissue, a 1-2 cm portion of the ileum was harvested, washed with PBS, and moved to a 24-well plate (Corning) filled with 200 μl RPMI (Gibco) with antibiotics and 10% fetal calf serum (Gibco). Ileum sections were incubated at 37 °C for 18 h, and then the supernatant was collected. The cytokine concentrations in the supernatant were determined using a cytokine bead array kit (BD Biosciences) following the manufacturer’s instructions. Levels of each cytokine (pg/ml) were calculated using standard curves provided in the kit.

### Tissue isolation

To obtain single-cell suspensions, the mLNs were processed through a 100 μm pore-size cell strainer by mechanical disruption using a buffer containing RPMI-1640 and 10% Fetal Calf Serum (FCS). Cells were centrifuged at 400 ×g for 10 mins at 4°C and resuspended in fluorescence-activated cell sorting (FACS) buffer (eBioscience) for staining. Cells from the spleen were prepared by grinding the spleen through a 70 μm cell strainer (corning). Splenocytes were centrifuged at 400 ×g for 10 mins at 4°C, and red blood cells (RBCs) were removed using RBC lysis buffer (Biolegend) for 2 mins on ice. Splenocytes were washed in sterile PBS and then resuspended in FACS buffer for staining. To isolate the cells from lamina propria, intestines were removed, flushed using sterile PBS, and opened longitudinally. The intestines were then cut into small pieces and rinsed thoroughly in HBSS buffer containing 5% FBS and 25 mM HEPES. Epithelial cells were removed with 20 mins incubations in extraction buffer containing HBSS with 15 mM HEPES, 5 mM EDTA, 10% FCS in a 37 °C shaking incubator. Tissue sections were then incubated in pre-warmed RPMI media containing 62 μg/ml Liberase TL (Sigma) and 50 μg/ml deoxyribonuclease I (Sigma), with continuous stirring at 37 °C for 40 mins in a shaking incubator. Following the digestion step, the tissue was passed through a 100 μm mesh filter (corning), and the resulting cells were further purified using a 40%/ 80% Percoll (GE Healthcare) gradient in RPMI-1640 (20 mins at 850 ×g at room temperature without acceleration/deceleration). Cells were then washed in HBSS buffer containing 5% FBS and 25 mM HEPES and re-suspended in FACS buffer.

### Flow cytometry and intracellular cytokine staining

For flow cytometry, single cell suspension from mLN, spleen, and colon lamina propria were incubated with anti-CD16/32 antibody (Biolegend) to block non-specific antibody binding to cells and then stained with Fixable live/dead viability dye (Thermo Fisher Scientific) for exclusion of dead cells. Cells were then stained with the following anti-mouse fluorophore-labeled antibodies (mAbs): CD3 (FITC, Invitrogen), CD4 (BUV 395, Invitrogen), CD45.1 (BV 605, Invitrogen), CD 45.2 (APC e780, Invitrogen), Vα2 (PE-Cy7, Invitrogen). For Foxp3 T staining, surface staining was performed, followed by overnight fixation with the Foxp3 transcription factor buffer set (Invitrogen). The monoclonal antibodies Foxp3 (PE, Invitrogen), Rorγt (APC, Invitrogen), and T-bet (BV786, BD Biosciences) were used. For intracellular staining, cells were stimulated with the cell stimulation cocktail (Invitrogen) for 4 h. Surface staining was performed, followed by fixation and permeabilization (BD Cytofix cytoperm Plus kit, BD Biosciences). The monoclonal antibody IFN-γ (PE, Biolegend) was used. Samples were processed in LSR Fortessa X-20 (BD Biosciences) and analyzed with FlowJo Software (FlowJo LLC) and data were compiled using Prism software (GraphPad).

### *In vivo* T cell conversion assay

Nursing CD45.2 C57BL6 dams with infected pups were given ova 20 g/l (Sigma) in drinking water for 14 days. One day after the start of dietary ova, mice were injected intraperitoneally with 5 × 10^5^ naive ova-specific T cells isolated from the lymph nodes of adult naïve CD45.1 Rag^−/−^ OT-II T cell receptor transgenic mice. CD4^+^ T cells were labeled with cell trace violet (Thermo Fisher Scientific). Following the transfer, tissue of interest was harvested at the indicated time points for cell isolation and analyzed by flow cytometry to evaluate the phenotype of the CD45.1^+^ OT-II T cells.

### Immune response functional assay

Nursing CD45.2 C57BL6 dams with infected pups were given ova 20 g/l in drinking water for 7 days or drinking water alone. Then, mice were immunized subcutaneously with an emulsion of 100 μg ova in complete Freund’s Adjuvant (CFA) (Sigma Aldrich). One week post-immunization, draining lymph nodes were isolated, and 1x 10^5^ cells per well were seeded in a 96-well plate and restimulated in vitro with 2 mg/ml ova. Supernatants were collected after 72 h, and IFN-γ levels were measured using a cytokine bead array kit (BD Biosciences) following the manufacturer’s instructions. Cytokine (pg/ml) was calculated using standard curves.

### ELISA of ova-specific IgG in serum

For measurement of ova-specific IgG2c in serum, high-binding ELISA 96-well plates (Corning) were coated with ova (2 μg/ml), overnight at 4 °C. Plates were washed three times with PBS/ 0.05% Tween 20 and blocked using 1% BSA prepared in PBS/ Tween 20. The serum was diluted 1: 10 in 0.1% BSA prepared in PBS/ Tween 20 and added to each well at 4 °C overnight. After washing, HRP-conjugated anti-mouse-IgG (diluted at 1: 10,000) prepared in 0.1% BSA prepared in PBS/ Tween 20 was added to the plate and incubated for 1 h at room temperature. The plates were washed five times with PBS/ 0.05% Tween 20. After a final wash, 100 μl of HRP substrate (BD OptEIA TMB Substrate Reagent) was added and the reaction was stopped by adding 50 μl of 2 molar sulfuric acid. Absorbance was read at 450 nm. Levels of anti-ova IgG2c were expressed in OD values.

### Quantification and statistical analysis

All analyses were done using GraphPad Prism 10.2.2 software. A normality test was performed to check if the data was distributed normally. For normally distributed data, an unpaired Student’s t-test was used, and for multiple comparison analyses, a one-way ANOVA was used for analysis. For non-normally distributed data, the Mann-Whitney test was used for comparison between two groups, and the Kruskal–Wallis test was used to compare the differences between groups. For *in vivo* experiments, each dot represents data from an individual mouse.

## ACKNOWLEDGMENT

We thank William Witola (University of Illinois at Urbana-Champaign) for providing the *C. parvum* oocyst. We acknowledge the core facilities-Digestive Disease Research core, Flow Cytometry & Fluorescence Activated Cell Sorting Core, Center for Genome Sciences & Systems Biology, Immunomonitoring Laboratory-Bursky Center for Human Immunology and Immunotherapy Programs, at Washington University School of Medicine in St. Louis. The Digestive Disease Research core center is supported by Grant #P30DK052574. The Center for Genome Sciences and Systems Biology is partially supported by NCI Cancer Center Support Grant #P30CA91842 to the Siteman Cancer Center and by ICTS/CTSA Grant# UL1TR002345 from the National Center for Research Resources (NCRR). This study was supported by grants from the NIH (AI171858 to L.D.S.).

---

**Supplementary Figure S1, related to Fig 1.**
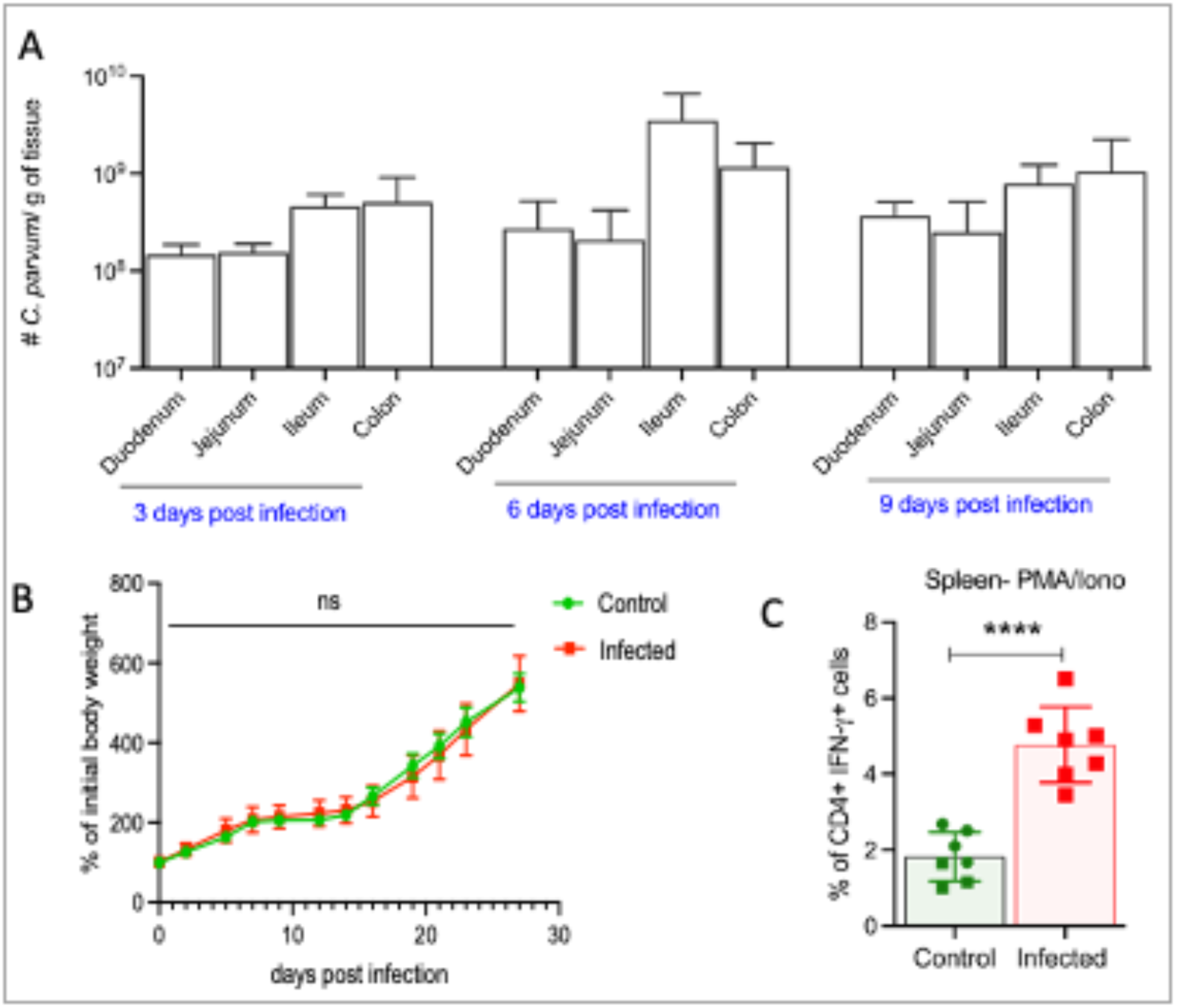
(A) The average number of *C. parvum* per g of tissue in 5-day old mice monitored at 3-, 6-, and 9-days post-infection by qPCR. (B) The percentage of initial body weight of 5-day-old control and infected mice monitored at the indicated days post-infection. (C) The frequency of CD4^+^IFN-γ^+^ T cells in spleen of 5-day old mice examined at 6 days post-infection obtained after activation with PMA/ionomycin for 3.5 hours and analyzed by flow cytometry. (A) Means ± SD from one independent experiment with n = 2-3 mice per group. (B) Means ± SD from one independent experiment with n = 6-9 mice per group. (C) Means ± SD from two independent experiments with n = 3-4 mice per group in each experiment. Statistical tests performed: (B) 2way ANOVA with Sidak’s multiple comparison test (C) Unpaired Student’s t-test, \*\*\*\**P* < 0.0001, ns, not significant.

**Supplementary Figure S2, related to Fig 2.**
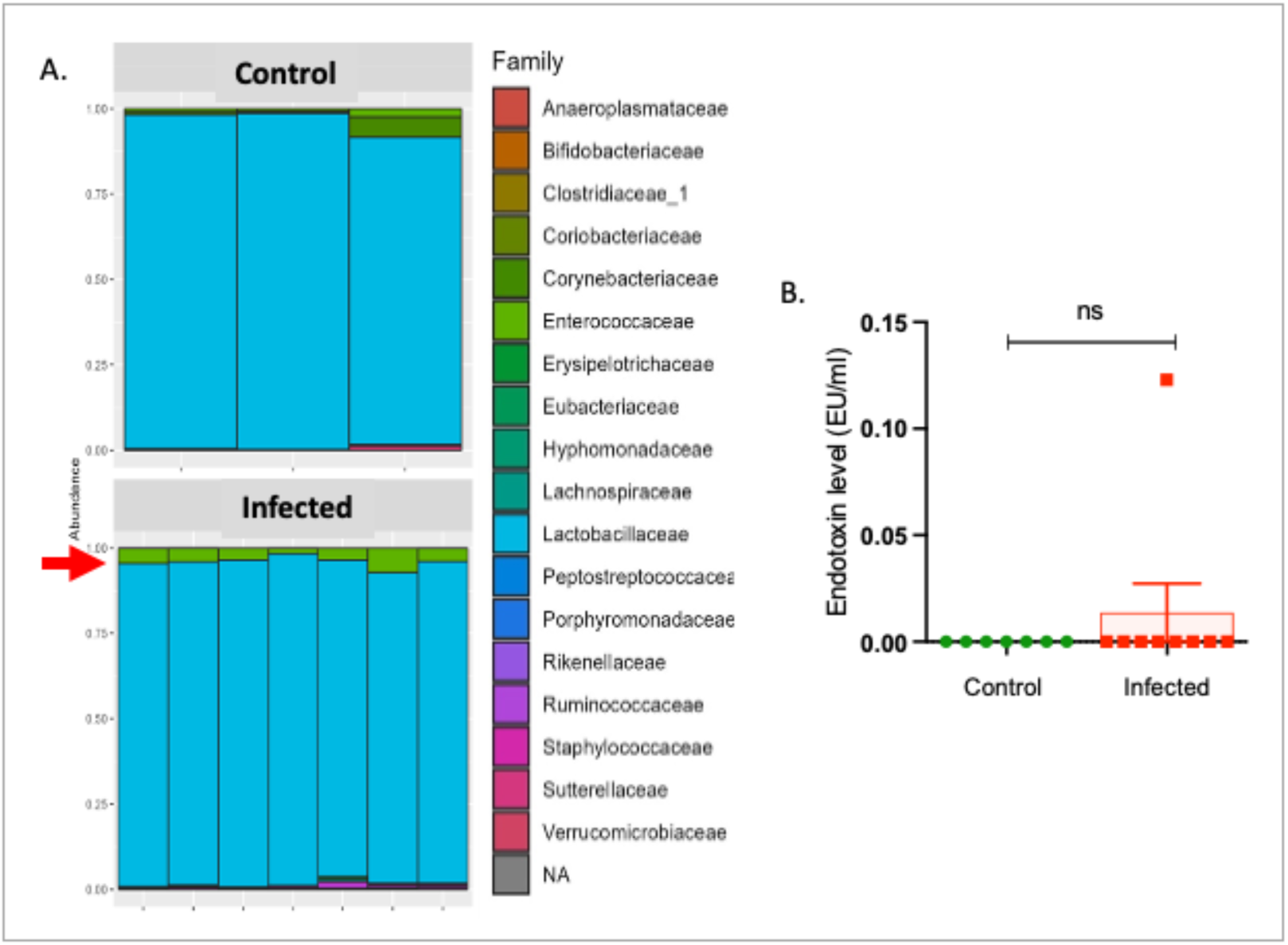
(A) Family level composition in 5-day old mice sampled at day 6 post-infection (B) The endotoxin levels in the serum of 5-day old mice measured at day 6 post-infection. Means ± SD from two independent experiments with n = 4-5 mice per group in each experiment. Unpaired Student’s t-test was used for comparison. ns, not significant.

**Supplementary Figure S3, related to Fig 3.**
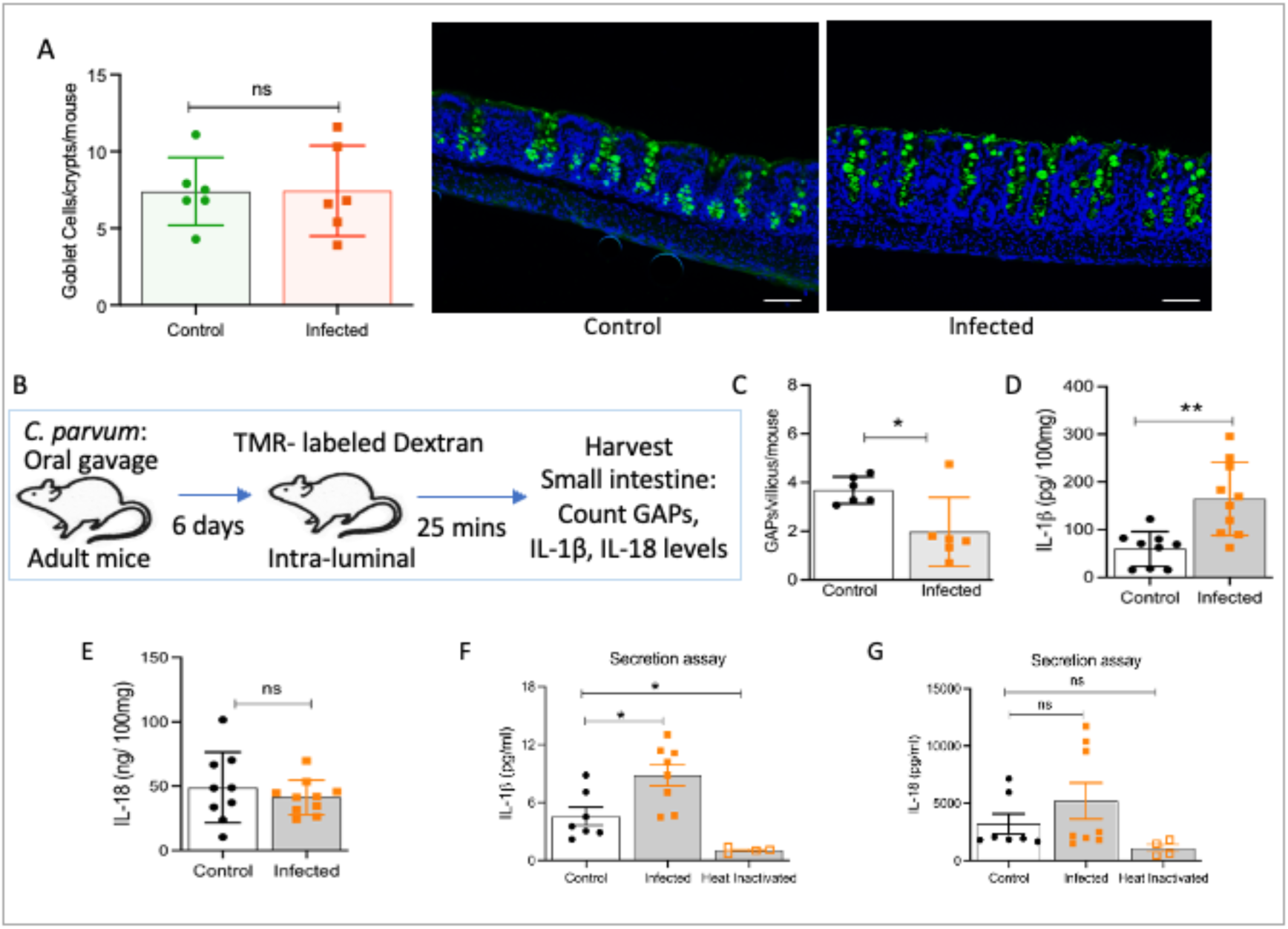
(A) Number of Goblet cells in the colon of 5-day old mice measured at day 6 post-infection, and their representative images, Scale bar, 50 µm. (B) Experimental design for counting GAPs and measuring levels of IL-1β and IL-18. (C) Frequency of GAPs in the small intestine of control and infected adult mice. Levels of (D) IL-1β, (E) IL-18 in the supernatant of epithelium scraping of adult control and infected mice by cytokine bead array. Levels of (F) IL-1β, (G) IL-18 in the supernatant of intestinal segment of adult control and infected mice incubated at 37 °C for 18 h by cytokine bead array. (A) Means ± SD from one independent experiment with n = 6 mice per group. (B-G) Means ± SD from two independent experiments with n = 3-4 mice per group in each experiment. Statistical tests performed: (A, C-E) Unpaired Student’s t-test was used for comparison. (F, G) One-way ANOVA with Sidak’s multiple-comparison test. \**P* < 0.05, \*\**P* < 0.01, ns, not significant.

**Supplementary Figure S4, related to Fig 4.**
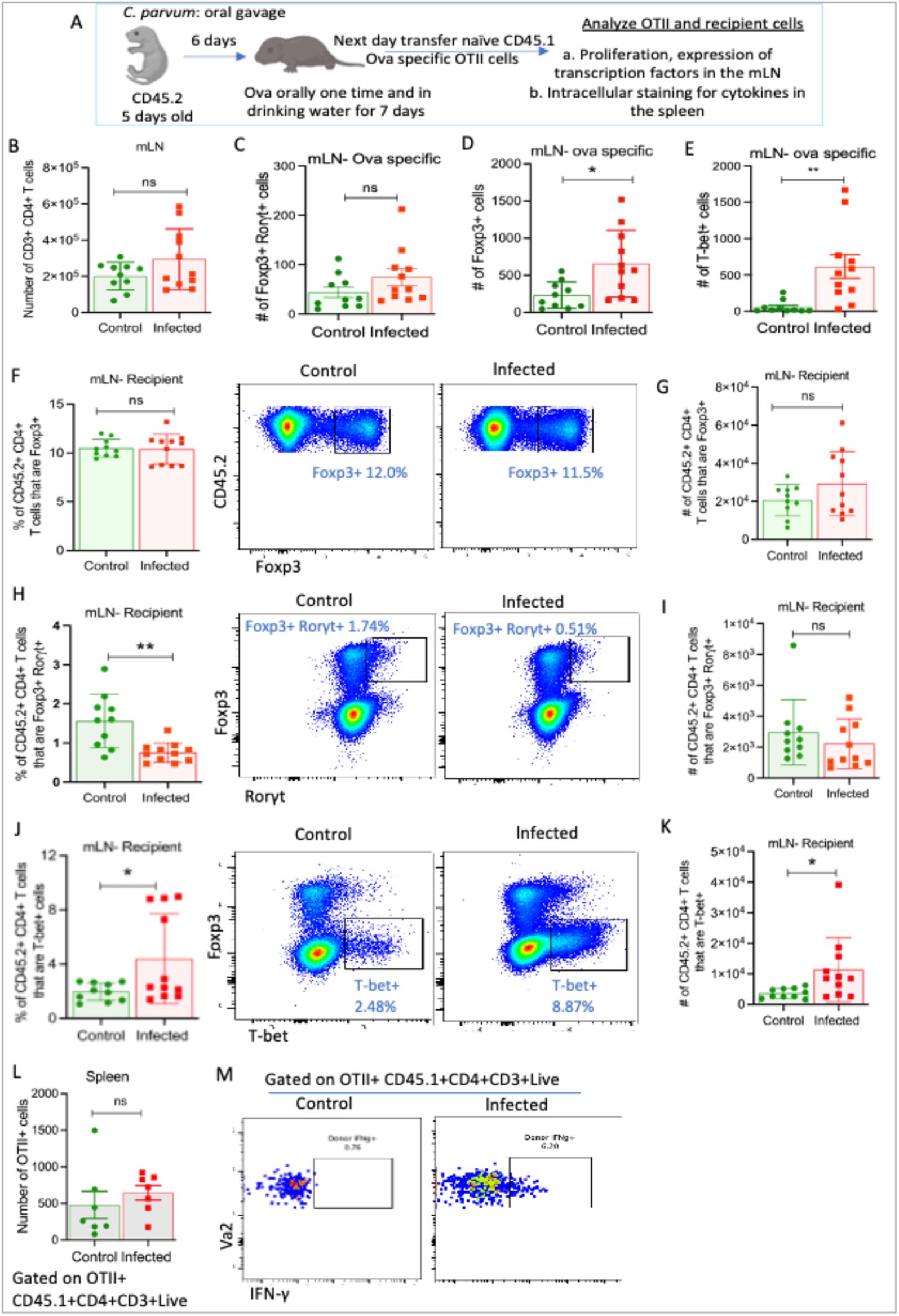
(A) Experimental design for monitoring CD4 T cells response to *C. parvum* infection in response to dietary antigens in the mLN. Number of (B) CD3^+^CD4^+^ T cells (C) ova-specific Foxp3^+^ cells (D) ova-specific Foxp3^+^Rorγt^+^ cells (E) ova-specific T-bet^+^ cells in the mLN of control and infected mice. (F-K) The frequency or number of different transcription factor-expressing recipient cells in the mLN of control and infected mice as shown. Cells are gated on CD45.2^+^ CD4^+^ CD3^+^Live cells (L) Number of OT-II cells in the spleen of control and infected mice. (M) Representative graphs for Fig 4H. (B-M) Means ± SD from two independent experiments, n= 3-4 mice per group in each experiment. Statistical tests performed: Unpaired Student’s t-test, \**P* < 0.05, \*\**P* < 0.01, ns, not significant.

**Supplementary Figure S5, related to Fig 5.**
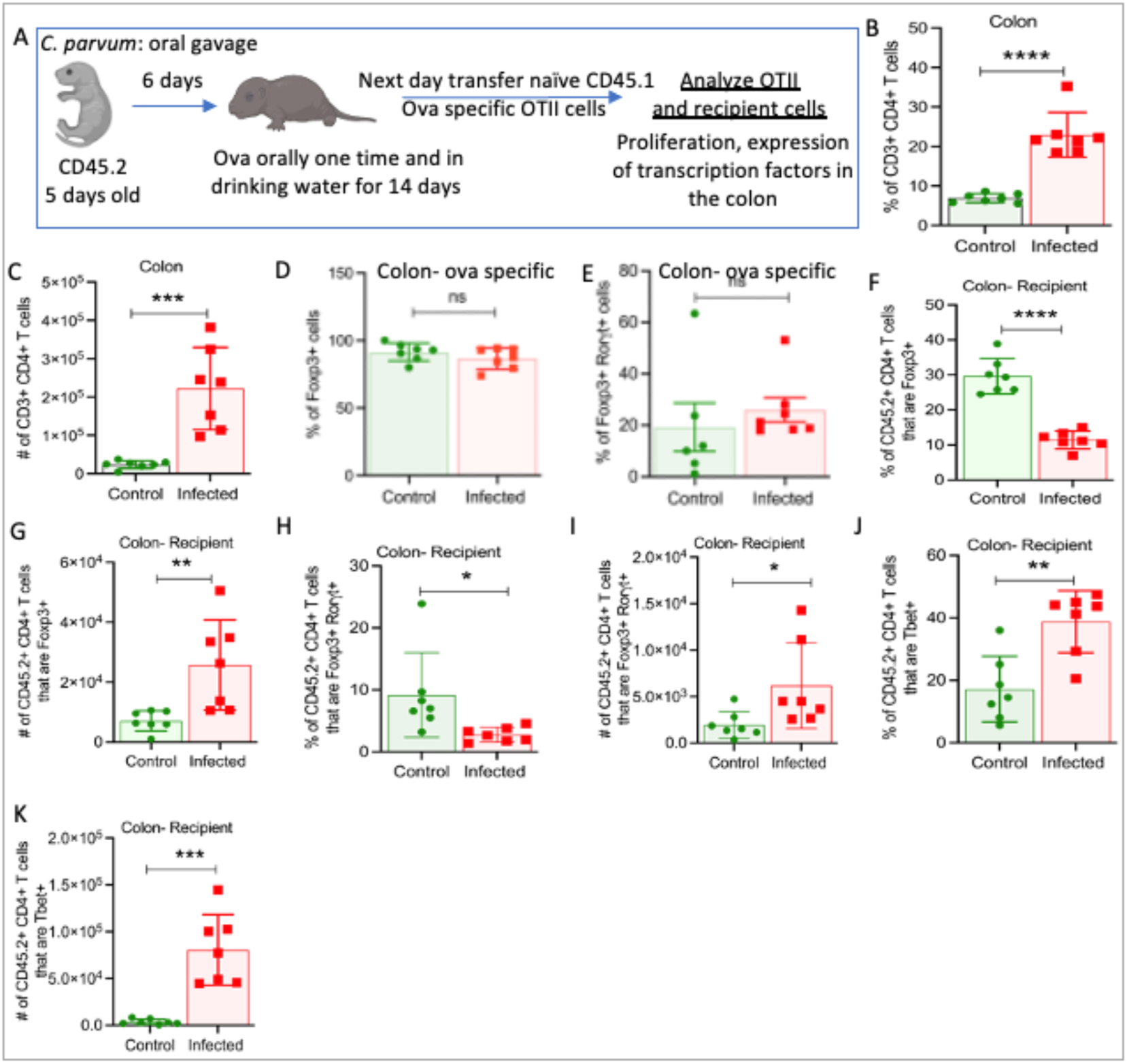
(A) Experimental design for monitoring CD4 T cells response to *C. parvum* infection in response to dietary antigens in the colon. (B) Frequency (C) Number of CD3^+^CD4^+^ T cells in the colon of control and infected mice. (D-K) The frequency or number of different transcription factor-expressing recipient cells in the colon of control and infected mice as shown. Cells are gated on CD45.2^+^ CD4^+^ CD3^+^Live cells (B-K) Means ± SD from two independent experiments, n= 3-4 mice per group in each experiment. (B-K) Unpaired Student’s t-test, One-way ANOVA with Sidak’s multiple-comparison test. \**P* < 0.05, \*\**P* < 0.01, \*\*\**P* < 0.001, \*\*\*\**P* < 0.0001.

**Table S1.**
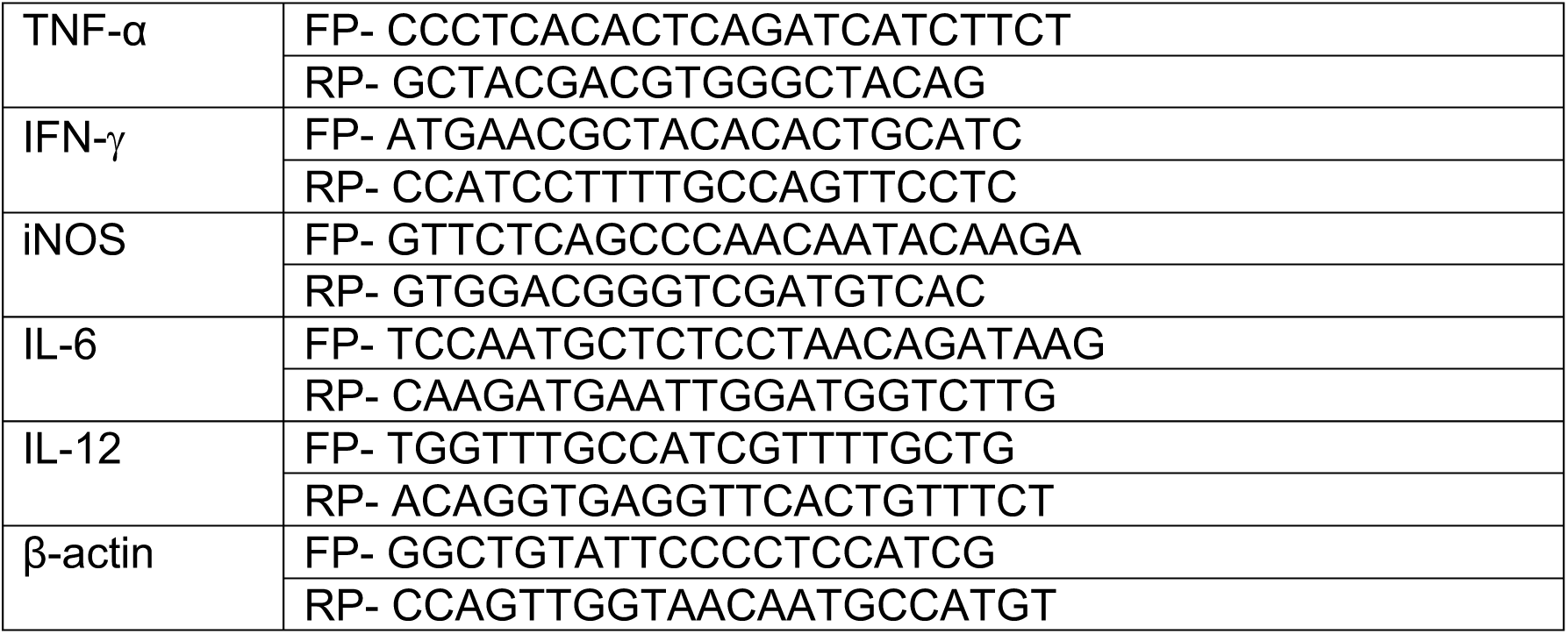
List of Primers for cytokine amplification. Related to Figure 1.

